# Comparing Methods for Mass Univariate Analyses of Human EEG: Empirical Data and Simulations

**DOI:** 10.1101/2025.11.19.689266

**Authors:** Anna-Lena Tebbe, Christian Panitz, Andreas Keil

**Affiliations:** University of Florida, Department of Psychology, Laboratory for Brain, Body, and Behavior; University of Bremen, Germany

**Keywords:** Multiple comparison, permutation tests, ERP, effect size, time frequency, Bayesian statistics, power

## Abstract

Electroencephalography (EEG) is a widely used method for investigating human brain dynamics. However, EEG analyses are frequently conducted with limited a priori knowledge regarding locations or latencies of meaningful statistical effects. This makes it difficult for researchers to form regions of interest (ROIs), which are then analyzed using traditional statistical models such as analysis of variance. In addition, exploratory studies, or studies interested in determining the exact temporal and spatial extent of a predicted effect may aim to examine many sensor locations and time points, often jointly. To address this, mass univariate analyses have become a valuable complement to ROI-based approaches. These methods attempt to correct for multiple comparisons while mitigating the risk of false positives and false negatives, thus enabling statistical inference in high-dimensional EEG data. Here, we review and evaluate different approaches for delineating spatial and temporal effect boundaries in three different datasets, focusing on within-subjects comparisons. Specifically, we focus on permutation-based approaches and their Bayesian alternatives to address condition differences in i) steady-state evoked responses, ii) event-related potentials, and iii) time-frequency data. Overall, simulation results indicate that cluster-based permutation tests provide a relatively liberal approach to correct for multiple comparisons across domains, with high sensitivity for detecting large effects. In contrast, the permutation-based *t*_max_ procedure yields the most conservative method across datasets. Bayesian approaches inherently are continuous in nature and thus strongly depend on the selection of thresholds for when support for a hypothesis is considered meaningful.

**Highlights:**

- Direct comparison of different mass univariate tools applied to real EEG data
- Variability in the number of datapoints showing statistical condition differences
- Mass univariate tools alleviate the arbitrary averaging of time windows and electrodes
- Caution is warranted when using Bayes factors for mass-univariate comparisons

## 1. Introduction

Electroencephalography (EEG) measures voltage changes, often across multiple spatial locations with millisecond precision. Especially since the advent of high-density electrode arrays with hundreds of channels, researchers confront datasets of high dimensionality: A typical data set may include hundreds or thousands of time points, recorded across dozens or hundreds of sensor sites, and spanning numerous frequency bands. When little a priori knowledge is available regarding the exact latency of a given statistical difference, its scalp distribution, or its frequency range, then analyzing the data with a mass univariate approach is particularly useful. Different methods for mass univariate analyses are available. However, it is less clear how to select from among these methods. They may differ in their sensitivity and specificity for detecting statistical and topographical differences, but there is a lack of systematic comparisons and quantitative analyses of these properties. The present study compares sensitivity to experimental effects across several mass univariate methods in real and simulated EEG responses such as event-related potentials (ERPs) and time-frequency data, across different effect sizes.

EEG waveforms represent neural mass activity, captured as voltage changes over time. Given the superposition of contributions from different brain sources and the biophysical properties of the brain (e.g., volume conduction), electrical activity is diffusely spread and recorded across multiple electrodes. In addition, the underlying cortical networks often exhibit synchronized activity over time, creating spatially and temporally dependent signals. Thus, a widely held assumption is that a meaningful statistical effect in the data should have some extent across the dimensions of interest, i.e., across 1) time, 2) frequency, and/or 3) spatial location, with true effects spanning multiple adjacent time points, neighboring electrodes, and frequency bands. As a results, researchers often reduce the dimensionality of the data and limit the number of statistical tests they perform. For example, to analyze ERPs, voltage may be averaged or integrated across an ROI containing time points and/or electrodes, resulting a in a low-dimensional dependent variable which is suitable for traditional statistical methods. How these specific electrodes and time points are selected has been subject to extensive debate, including questions regarding replicability and robustness. Regions of interest are often determined a-priori, based on existing literature or theory, or selected based on visualization and screening of the data. Condition differences are then typically contrasted using parametric tests, such as t-tests or ANOVAs.

Challenges of the ROI-based approach include that many ERPs do not have a clear onset, and that their topographical distribution and temporal extent tends to be diffuse and highly variable in different studies (Clayson et al., 2013; Groppe et al., 2011a; Intriligator & Polich, 1995; Kiesel et al., 2008). Furthermore, a researcher’s expectations and decisions along the preprocessing and analysis stream – e.g., selecting specific time points, frequency ranges, or sensor sites (Dien & Santuzzi, 2004; Groppe et al., 2011a) – can affect and bias the results (Luck & Gaspelin, 2017; Steegen et al., 2016; Wicherts et al., 2016). In addition, with this approach the wealth of information embedded in the EEG data is not fully leveraged: While time-window or sensor-group averages can indicate the presence of an effect, they provide limited insight into its precise timing and spatial distribution (Groppe et al., 2011a).

On the other hand, analyzing differences with univariate statistical methods, at all possible electrode by timepoint combinations, would also drastically increase the probability of committing false statistical inferences in cases where researchers apply null-hypothesis testing (NHT) methods: When many individual tests are conducted, under the null hypothesis, the false positive error rate (alpha) inflates, so that the chance of at least one false positive exceeds the commonly used value of alpha = .05 (multiple comparisons problem). To mitigate this problem, corrections for multiple comparisons have been applied. A well-known exemplar to control for false positives is the Bonferroni correction, which adjusts the family wise error rate (Type I error) by dividing the critical *p*-value by as many comparisons as have been conducted. Yet, given a total of 2000 comparisons that might have to be corrected for, an exemplary *p*-value of .05 would shrink to a critical value of .0000025, rendering this an overly conservative, insensitive, and likely invalid option. In addition, Bonferroni correction assumes that each of the tests is independent, which is not the case with EEG data, as adjacent values (e.g., timepoints or electrodes) are highly correlated. Alternative methods for addressing the multiple comparison problem have been *permutation approaches,* for example by examining temporal and spatial clusters (Maris & Oostenveld, 2007; Meyer et al., 2021) or using other characteristics based on the *t*-statistic (Blair & Karniski, 1993). Specifically, a more conservative critical *t*-score is derived via permutations of the empirical data, and condition differences above this permutation-based threshold are considered significant. In addition, *Bayesian statistics* can complement or replace the null hypothesis testing (NHT) approach by continuously quantifying evidence for or against a hypothesis, without relying on p-values or interpreting results in terms of the probability of observing data under a null hypothesis. Characteristics of these methods are elaborated in more detail in the following paragraphs.

### 1.1. Cluster-Based Permutation Test

Cluster-based permutation can be used for assessing the statistical significance in a variety of psychophysiological research questions, where the hypothesis about the existence of an effect (e.g., a difference in response amplitude between two experimental conditions), but is agnostic to the temporal characteristics (i.e., the precise timepoints of its emergence and duration) and spatial extent (i.e., the exact number of neighboring electrodes that are considered part of the cluster).

The procedure identifies statistically meaningful clusters of adjacent time points, electrodes, frequencies or other dimensions, in which adjacency is possible by testing the significance of the observed cluster against bootstrapped permutations of the same data when condition labels are randomly swapped within subjects or subjects are (pseudo-)randomly assigned to groups. The current manuscript focuses on within-subject effects. Under the assumption of exchangeability, permutation procedures sample the data via repeated random shuffling of condition labels, yielding a null distribution of cluster-mass statistics, e.g., of *t*-scores (Good, 2005; Groppe et al., 2011a, 2011b).

Cluster-based permutation tests consist of the following procedural steps:

1. First, a sample statistic, such as a *t*-score, is calculated comparing condition A and condition B (statistic *_observed_*) for each variable, e.g., each time point, frequency band, and electrode of interest. From all resulting statistics, only those that exceed a predefined threshold (*cluster-forming threshold*, e.g., a *t*-score corresponding to an uncorrected *p*-value of .05) are considered.
2. These statistics are grouped into clusters based on their proximity in time, space, or frequency. Here, an a-priori definition of *neighborhood* is needed, e.g., a certain spatial distance of electrodes on the scalp (see Maris & Oostenveld, 2007).
3. A cluster-mass statistic is derived (statistic *_sum_*), e.g., by taking the sum of any cluster’s t-scores or number of members in any cluster (Bullmore et al., 1999).
4. The last step is the *permutation*, which is used to approximate a null distribution against which the observed cluster-mass statistic is compared. To generate this null distribution, the condition labels are randomly swapped, and new cluster-mass statistics are calculated as described above. Then, the cluster with the greatest mass statistic is identified and its value saved. This process is repeated many times, and the saved maximum cluster mass statistics form a reference distribution under the null hypothesis, i.e., that there is no difference between conditions. The statistical significance of a cluster is determined by the probability of observing the actual cluster-mass statistic (statistic *_observed_*), given the permutation-based null distribution. For example, a two-sided *p*-value of .05 is tested by comparing against the 97.5^th^ percentile of the permutation-based null distribution (Sassenhagen & Draschkow, 2019; for a tutorial, see Maris & Oostenveld, 2007; Groppe et al., 2011a).

For most cases, performing a full permutation of all possible combinations is computationally intractable. Instead, approximations are used by repeatedly calculating a sample statistic (e.g., *t*-score or *F*-value) on many permutations of the data (1,000–10,000 repetitions), yielding results similar to a full permutation (Blair & Karniski, 1993; Groppe et al., 2011a; Manly, 2018; Sassenhagen & Draschkow, 2019). Previous work has urged researchers to use caution with regard to the interpretation of clusters, particularly their precise time window including on- and offset of significant effects or their specific ‘cluster members’ (Sassenhagen & Draschkow, 2019). Because *p*-values are specific to the cluster-level statistic, they cannot be transferred or extended to each timepoint or electrode within a cluster. To accurately define effect boundaries while controlling error rates, other statistical tests can be applied (e.g., Tukey’s HSD or jack-knifing, see (Groppe et al., 2011a; Miller et al., 2009; Sassenhagen & Draschkow, 2019; Smulders, 2010).

### 1.2. t_max_ Permutation

Another permutation-based approach was initially described by Blair & Karniski (1993). It also assesses statistical significance based on permutation-based thresholding of empirical, mass-univariate, t-scores, F-scores, or other test statistics. It is applicable to one-sample or repeated measures and follows a similar approach as the cluster-based permutation tests described above. In contrast to cluster-based testing however, the maximum test statistic across all data points from each iteration of the permutation procedure enters the test distribution without taking clusters or neighboring electrodes into account. Paralleling the cluster-based permutation test, an approximation of all possible combinations of *t*-scores is calculated by repeatedly exchanging condition labels to form the reference distribution (e.g., the *t*_max_ or the (absolute) sum of all *t*-scores: *t*_sum_ or |*t* _sum_|). Each individual test score from the mass-univariate test array (e.g. a topographical map or time series) can then be compared against a suitable threshold based on a percentile of the t_max_ or t_sum_ distribution. Similarly to cluster-based permutation tests described above, there is no need to estimate the correlation or distribution of the underlying data.

### 1.3. Bayesian approaches

Bayesian statistics represent an approach to data analysis and parameter estimation based on Bayes’ Theorem, which allows researchers to quantify the evidence for or against a given hypothesis. Before collecting data and testing for a (statistical) relation between variables, there might be assumptions and preexisting information about this relationship (priors). This knowledge about the values for a given model parameter can be directly incorporated into the modelling process by a prior distribution. The observed evidence is then used to update these distributions, i.e., priors are combined with the likelihood of the data to form the posterior distribution, which provides the estimate for each model parameter. In contrast to frequentist approaches, Bayesian statistics allow direct probability statements about parameters while updating beliefs as new data become available in an iterative process, and do not face the multiple comparison problem to the extent that frequentist statistics do (e.g., Gelman et al., 2012). This difference arises because Bayesian statistics do not rely on *p*-values or null hypothesis significance testing, which make multiple comparison corrections necessary. However, if many comparisons are computed, priors should reflect reasonable constraints to avoid overfitting. Careful modeling and interpretative caution are important, especially when making many inferences.

It is important to note that Bayes Factors quantify the relative evidence for one statistical model over another model and thus do not indicate statistical significance as the *p*-value, but rather reflect a continuous measure of evidence for or against an effect. Labels such as “moderate” or “strong” evidence are often used to aid the interpretation (Lee & Wagenmakers, 2014; Jeffrey 1961).

Given these different approaches, the question arises if and how these methods vary in detecting differences in EEG responses between experimental conditions. It is challenging to determine if a procedure that performs well on one dataset will be equally effective for others. Similarly, it is unclear how increasing effect sizes influence the observed latency and topographical differences detected by mass univariate approaches. However, no study has contrasted the topographical and statistical implications of different methodologies for addressing the multiple comparison problem for both real and simulated EEG data. Here, we extend prior work by a series of simulations to review and compare different procedures to correct for the multiple comparison problem by examining their sensitivity and topographical effect distributions in real and simulated EEG datasets over varying effect sizes.

## 2. Methods

To compare different mass univariate approaches, we ran simulations on three example datasets in the time and frequency domain, applying permutation approaches and Bayesian t-tests.

All participants had normal or corrected-to-normal vision and reported a negative personal and family history of photic epilepsy. Written informed consent was obtained from all participants. Procedures were approved by the institutional review board of the University of Leipzig (dataset 1) and the University of Florida (dataset 2). The third dataset consists of simulated EEG data.

### 2.1. First Dataset: Steady-State Visually Evoked Responses

The first dataset is from a study examining steady-state visually evoked responses (SSVEP) in a perspective taking paradigm (Tebbe et al., 2024). SSVEPs are oscillatory neural responses elicited by a visual stimulus that is presented at a constant frequency (see Wieser et al., 2016). In the experiment, participants watched videos in which a person observed flickering objects (4 Hz) moving behind an occluder or inside a tunnel, evoking SSVEPs at 4 Hz. In each trial, the person followed the object with her gaze as it moved across a table and then slowly disappeared behind an occluder or into a tunnel and remained occluded until the end of the trial. Once the object moved behind the occluder, the participants’ view to the object was blocked, whereas the person in the video continued to have visual access to the flickering object. In contrast, both the participants’ and the person’s visual access was blocked in the Tunnel condition. The total duration of the videos was 9.5 sec.

The study reports EEG data of *N* = 40 subjects (15 female, 25 male; age *M* = 26.05 years, *SD* = 3.66). Two additional subjects participated but were excluded from further analysis due to more than 50% EEG trials with artifacts. Continuous EEG was recorded using 28 active electrodes placed in an electrode cap (Easycap GmbH, Herrsching, Germany) according to the International 10–20 System. Electrode impedances were kept below 30 kΩ. Data were digitalized with a sampling rate of 500 Hz with Cz as reference and offline preprocessed using Brain Vision Analyzer (Version 2.1.2.327) and MATLAB (2023a; Mathworks, Inc., Natick, MA, USA). Epochs with major artifacts were identified by visual inspection. Further segments with artifacts were identified using an automatic artefact correction with the following criteria: 1) a maximal allowed voltage step of 50 μV per ms; 2) a maximal difference in amplitude (maximum-minimum) of 300 μV in a time interval of 300 ms; 3) a minimum or maximum amplitude of ±500 μV; 4) and the lowest allowed activity in time intervals of 100 ms set to 0.5 μV. Data was band-pass filtered to 1-20 Hz with an IIR zero phase Butterworth filter. Eye Movements were automatically corrected using the Ocular Correction Independent Component Analysis (ICA) implemented in the Brain Vision Analyzer Software.

For the SSVEP analysis, artifact-free trials were first averaged for the Occluder and Tunnel conditions separately. Data were then bandpass filtered with a 10^th^ order zero phase Butterworth filter (half-width of 1 Hz) around the stimulation frequency (4 Hz). The time-varying amplitude of the SSVEP response at 4 Hz was extracted using the Hilbert transform (for details, see Tebbe et al., 2024). To eliminate offset differences between conditions, the mean amplitude of a time segment between 4500-6234 ms was subtracted as baseline, in which the object was fully visible. For the present study, the resulting baseline-corrected SSVEP responses were shortened to a segment from 6234-7500 ms post onset, corresponding to the time window of the object’s disappearance, i.e., starting with the first frame of when the object began to move behind the occluder or into the tunnel until its full occlusion. We hypothesized a prolonged 4 Hz response when the agent continued to see the flickering object in the occluder condition compared to when her visual access to the object was blocked (tunnel condition).

### 2.2. Second Dataset: Event-Related Potentials

The second dataset reports ERP data from *N* = 18 students of the University of Florida, (8 female, 10 male; age *M* = 20.22 years, *SD* = 1.44). Participants were presented flickering stimuli of faces and scenes (see Tebbe et al., 2021). In a subsequent recognition task, participants were presented with a total of 144 pictures (trials), of which 72 pictures were targets (previously seen) and 72 trials were distractors (novel pictures). Participants were asked to classify stimuli into ‘previously seen’ and ‘novel’ pictures via mouse button press. Only correct responses were analyzed and were categorized as i) hits (i.e., correctly identified as previously seen) or ii) correct rejections (i.e., correctly identified novel picture). Previous research has reported a more positive waveform for hits in comparison to potentials evoked by correct rejection (Curran & Friedman, 2004; Rugg & Curran, 2007), typically recorded over centro-parietal areas in an approximate time window of 500-800 ms (referred to as old-new effect; Weymar et al., 2011).

EEG was continuously recorded from 129 electrodes with a HydroCel Electrical Geodesics (EGI) system at a sampling rate of 500 Hz with vertex sensor (Cz) as the reference electrode. All electrode impedances were below 40 kΩ. EEG data were filtered offline with a Butterworth lowpass-filter at 40 Hz and a Butterworth highpass-filter at 0.1 Hz, implemented in the EMEGS (ElectroMagnetoEncephalograph) toolbox for Matlab (Peyk et al., 2011). Trial epochs were extracted from 100 ms pre and 800 ms post image onset. Trials with artifacts were identified using distributions of statistical parameters of the epochs (i.e., absolute value, standard deviation, maximum of the differences) for each trial and channel using the artifact rejection method proposed by Junghöfer and colleagues (2000). An average (± SD) of 100.78 ± 22.23 of 144 trials across individuals were retained. The number of channels excluded ranged from 1 to 11 (*M* = 4.56, *SD* = 3.32).

### 2.3. Third Dataset: Simulated Alpha Activity

For the third dataset, EEG data were simulated as Brownian noise (i.e., the cumulative sum of stochastic, white noise time series) with an epoch length of 3000 ms for 50 subjects across two experimental conditions, referred to as ‘blocking’ and ‘no blocking’. The simulation designated the initial 1200 ms of each EEG trial epoch as the baseline segment, followed by a simulated stimulus onset, and 1800 ms of post-stimulus time. For each of the two simulated conditions, 60 trials of Brownian noise were generated per subject for a single electrode. To simulate event-related changes in alpha activity in the blocking condition, a 10 Hz sine wave time series with a duration of 800 ms (8 full cycles of simulated alpha activity) was generated. This simulated alpha-band signal was systematically manipulated in its amplitude (see below) and added to the Brownian noise of the baseline segment of each trial in the blocking condition, beginning at 400 ms and ending at the simulated stimulus onset (1200 ms into each segment). After baseline adjustment of the evolutionary spectrum as described below, this manipulation leads to a selective, sustained decrease in alpha power relative to baseline in the blocking condition, scaled by the magnitude of the simulated 10 Hz sine wave.

To analyze temporal dynamics of oscillatory activity in the simulated alpha data, we used complex Morlet wavelets. A Morlet parameter of 10 was defined, resulting in a frequency smearing (σ_f_) of 1 Hz and a temporal smearing (σ_t_) of 159 ms at the frequency of interest, 10 Hz. The convolution of the wavelets with the simulated data was performed at the single trial level, for frequencies ranging from 1.333 Hz to 19.333 Hz in steps of .333 Hz, providing broad coverage of the alpha band and lower frequency bands. Condition averages were then calculated for each participant and condition separately, by averaging the single-trial time-frequency representations. The resulting averaged evolutionary spectra were normalized to the mean of a baseline segment (500–1000 ms relative to the simulated stimulus onset) by calculating percent change from baseline. Baseline corrected time-varying spectral amplitudes (in percentage change) were then used for further statistical analyses. In the present study, alpha power was analyzed as the power captured by wavelets with a center frequency at 10 Hz.

### 2.4. Statistical Analyses

Statistical analyses were performed in Matlab (R2024a).

Each of the datasets and methods underwent a three-step procedure. First, different effect sizes were simulated by manipulating the condition differences in the time window of the effect in the original dataset (dataset 1: 6234-7500 ms; dataset 2: 500-800 ms; dataset 3: 1200 ms pre-stimulus). That is, one of the two conditions was scaled by multiplication such that condition differences increased linearly from a maximum Cohen’s d of *d* = 0 to a maximum of *d* = 2 for all datasets (i.e., maximum *d* across the timepoints of the effect, electrodes, and – if applicable – frequency bands). Note that the exact effect size varied across electrodes (or frequency bins), so that the maximum Cohen’s d was used to reflect the strongest observed effect. In a second step, we obtained critical *t*-scores through permutation for each effect size separately. To this end, conditions labels were randomly shuffled, and *t*-score matrices (dataset 1 and 2: electrode by time point; dataset 3: time point by frequency bin) were recalculated 5000 times. For Bayesian t-tests, we used a critical threshold of BF_10_ ≥ 3 and BF_10_ ≥10. As a last step, when *t*-scores or BFs were exceeding the respective threshold, we extracted the number of datapoints (timepoints by electrode pairs or frequency by electrode pairs) for each effect size separately to contrast the sensitivity of each of the methods. The procedure was applied for each of the three datasets across the range of simulated effect sizes (i.e., Cohen’s *d* varying from 0 to 2, see Fig. 1).

**Figure 1.**
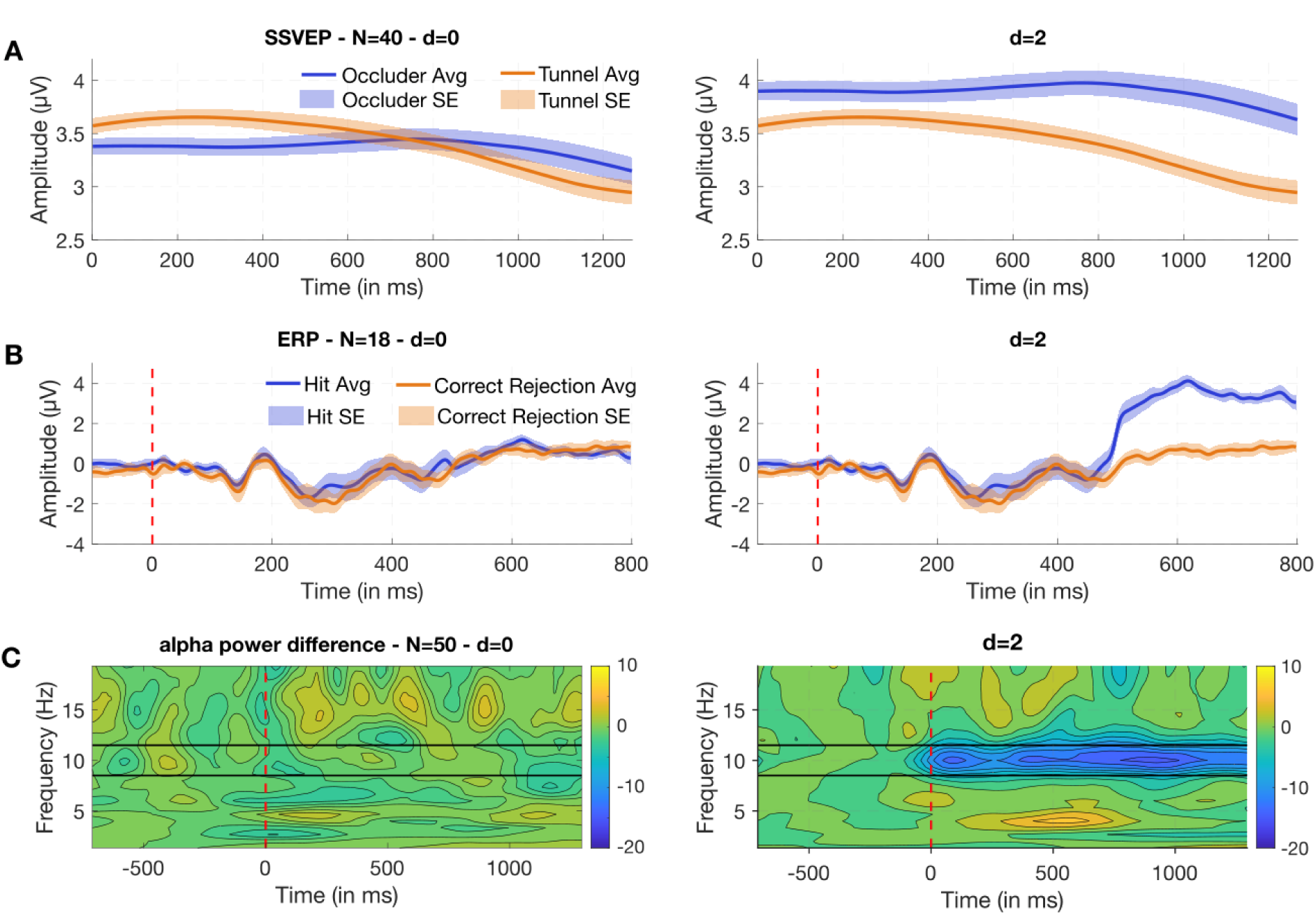
Illustration of the simulated effect sizes across the three datasets. Simulated condition differences ranged from no effect (*d* = 0, left column) to a large effect with a maximum Cohen’s *d* of 2 (right column). (**A**) Simulation results for the first dataset, reporting SSVEP responses over visual cortex (*N* = 40; electrode Oz). Participants showed a higher SSVEP amplitude for the Occluder (blue) compared to the Tunnel (orange) condition. (**B**) Simulation results for the second dataset, showing ERP waveform amplitude in a recognition task, indicating an old-new effect over parieto-central sensor sites at around 500-800 ms (*N* = 18; sensor 55) with greater responses to previously presented pictures (hits, blue) compared to correctly rejected stimuli (orange). Shading indicates the standard error. (**C**) Time-frequency differences between two conditions in the third dataset for *N* = 50 participants. Alpha power was simulated at the 10 Hz frequency. A reduction in alpha power was simulated in the time window of -800 ms until 0 ms relative to a simulated stimulus onset (0 ms, red dashed line).

For the *cluster-based permutation analyses*, a cluster’s timepoint and electrode members were defined with a custom image processing algorithm to find connected components over time and space. This resulted in approximate average of 3.5 neighbors for each electrode for the 26-channel layout (dataset 1) and 5.6 for the EGI layout (dataset 2). For the simulated alpha data (dataset 3), cluster members were identified by detecting connected components across time and adjacent frequency bins.

For *forming* a cluster, a threshold of *p_crit_* = .05 or *p_crit_* = .01 of the student *t*-distribution was used (*cluster-forming threshold*), corresponding to a critical *t*-score of ± 2.07 and ± 2.81 for the SSVEP data (dataset 1, *N* = 40); a *t*-score of ± 2.1 and ± 2.93 for the ERP recordings (dataset 2, *N* = 18). For the simulated alpha data (dataset 3, *N* = 50), this corresponded to a critical *t*-score of ±2.06 and ±2.8 for a threshold of *p_crit_* = .05 or *p_crit_* = .01, respectively. Values above these positive and below the negative thresholds were considered as part of a cluster when they were connected in time and electrode space, or in time and frequency space, respectively.

For a *cluster-forming threshold* at *p* = .05, the resulting critical threshold (i.e., the sum of all *t*-scores within a cluster) at a maximum Cohen’s *d* of *d* = 0 computed via permutation were 3687.22 at *p_crit_* = .05 (i.e., the 97.5^th^ percentile of the distribution) and 5125.72 for a *p*-value of *p_crit_* = .01 (i.e., the 99^th^ percentile) for the SSVEP data (dataset 1). For the ERP recordings (dataset 2), this resulted in a *t*-score sum of 5285.32 at *p_crit_* = .05 and 8801.02 at *p_crit_* = .01. For the alpha simulations (dataset 3), this resulted in a sum of -1003.31 at *p_crit_* = .05 and -1268.03 at *p_crit_* = .01.

For a *cluster-forming threshold* at *p* = .01, the resulting critical threshold following permutation at max. Cohen’s *d* = 0 were 1284.35 and 1987.41 for the SSVEP data (critical threshold of *p_crit_* = .05 and *p_crit_* = .01; dataset 1); a threshold of 887.81 and 8304.90 at *p_crit_* = .05 and *p_crit_* = .01 for the ERP recordings (dataset 2); and a threshold of -505.28 at *p_crit_* = .05 and -657.42 at *p_crit_* = .01 for the alpha simulations (dataset 3).

### t_max_ Permutation

For the *t*_max_ thresholds, we extracted the minimum and maximum *t*-scores in each permutation to form two reference distributions for each time point (*t*_max_ and *t*_min_ distributions). The i) 2.5^th^ and 97.5^th^ percentile and ii) 1^st^ and 99^th^ percentile of these distributions were used as critical thresholds, corresponding to an undirected *t*-test with a *p_crit_*-value of .05 and .01, respectively. Values more extreme than these thresholds were considered statistically significant. These *t*-scores corresponded to a maximum *t*-score of 3.97 and 4.46 for the SSVEP responses (*p_crit_* = .05 and *p_crit_* = .01; dataset 1), a maximum *t*-score of 5.31 (*p_crit_* = .05) and 6.68 (*p_crit_* = .01) for the ERP recordings (dataset 2), and a minimum *t*-score of -4.39 (*p_crit_* = .05) and -4.91 (*p_crit_* = .01) for the alpha simulations (dataset 3) when no condition differences were present (max. Cohen’s *d* = 0).

### Bayesian t-test

For *Bayesian* analysis, we contrasted each Bayes factor (BF_10_) within each dimension (time point, electrode, frequency bin) with the BF criterion set at BF_10_ ≥ 3 and BF_10_ ≥ 10. For each effect size, we ran Bayesian t-tests comparing all timepoints and electrodes for the SSVEP and ERP data (dataset 1 and 2) and all timepoints and frequency bins for the simulated alpha data (dataset 3). We used default priors for all models, i.e., an equal, uninformed prior probability using the bayesfactor toolbox for Matlab (Krekelberg, 2019). Although Bayes Factors quantify the relative evidence rather than providing dichotomous indicators of statistical significance, we here used two thresholds to allow for comparisons with other mass univariate tools. Following conventions, we considered a BF_10_ between 3 and 10 as moderate evidence, between 10 and 30 as strong evidence for the given hypothesis H1 (Lee & Wagenmakers, 2014). Accordingly, a BF_10_ between 1/3 and 1/10 was considered moderate evidence, between 1/10 and 1/30 as strong evidence in favor of H0. BFs between 1/3 and 3 were considered inconclusive.

## 3. Results

### 3.1. First dataset: Steady-State Visually Evoked Potentials

The simulation results are visualized in Figure 1, with datapoints plotted against the maximum Cohen’s *d* computed across all electrodes and time points for the different statistical approaches. As expected, higher thresholds for forming a cluster or a higher *t*-score threshold resulted in fewer datapoints surpassing the threshold after permutation testing. Similarly, a more conservative Bayes Factor threshold of BF_10_ ≥ 10 resulted in fewer datapoints exceeding the threshold compared to a BF_10_ ≥ 3. Consistently, the maximum number of electrodes exceeding the threshold increased with larger effect sizes. Further, the figures illustrate how stricter thresholds, regardless of whether a Bayesian or frequentist approach is used, reduced the number of datapoints and electrodes identified as indicating condition differences.

#### Cluster-based permutation

The cluster-based procedure did not indicate significant condition differences before a small to medium effect was simulated in the data. Specifically, for dataset 1, the first cluster was identified around a Cohen’s *d* of *d* = .2, see Fig. 1. The number of electrodes of the cluster gradually increased from below 3 (∼10%) to all electrodes (total of 26) at *d* = 1.4 for both cluster-forming thresholds of *p_crit_* = .05 and *p_crit_* = .01. As expected, with a more conservative cluster-forming threshold of *p_crit_* = .01, less datapoints and electrodes were detected. Overall, condition differences were comparable to a threshold set at BF_10_ ≥ 10. Compared to other techniques, the cluster-based permutation identified most electrodes as exceeding a meaningful threshold, with the steepest increase in electrodes and datapoints compared to the Bayesian t-test and the *t*_max_ procedure. Datapoints identified when using the 97.5^th^ percentile of the null distribution were highly similar to the 99^th^ percentile, indicating minimal difference between the two cut-offs.

#### T_max_ procedure

The *t*_max_ correction yielded the least significant datapoints overall and proved to be the most conservative technique compared to the other methods applied, also see Fig. 2. Both, the cluster-based and the *t*_max_ permutation identified no datapoints or electrodes below small effects (*d* = .2), however, the number of identified datapoints and electrodes using the *t*_max_ permutation was much lower compared to the cluster-based permutation. The overlap between datapoints (timepoints by electrodes) identified as significant using the cluster-based compared to the *t*_max_ permutation at Cohen’s *d* of *d* = .5 (medium effect) and *d* = .8 (large effect) was around 10%. The overlap in number of electrodes was between 10-30%. When Cohen’s *d* reached 1.1, the *t*_max_ procedure identified about 60% of the electrodes at a critical *p*-value of .05, compared to those detected with the cluster-based permutation and the Bayesian t-test with a threshold of BF_10_ ≥ 3 (65% overlap with a threshold of BF_10_ ≥ 10). Similarly, when applying a more conservative threshold of *p_crit_* = .01, the proportion of identified electrodes reached about half out of all electrodes at a Cohen’s d of *d* = 1.3, corresponding to above 90% of the electrodes identified when applying the cluster-based permutation or Bayesian approach.

**Figure 2.**
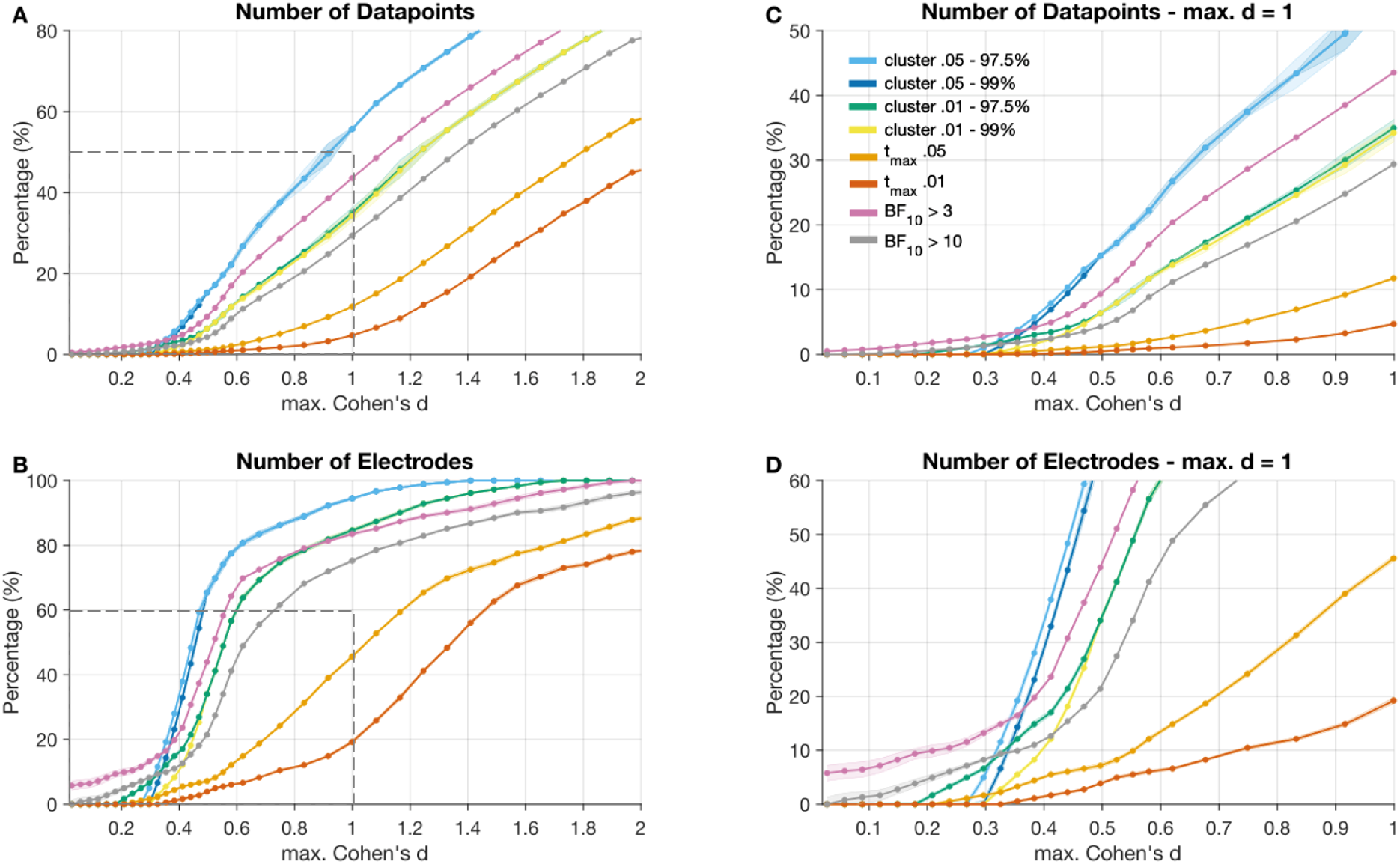
Results from the first dataset showing datapoints and electrodes that exceeded the predefined threshold for the different mass univariate approaches across simulated effect sizes for the SSVEP data. (**A**) Percentage of datapoints and electrodes exceeding the critical thresholds defined by the different statistical approaches: cluster-based permutation using different cluster-forming thresholds (*p_crit_* = .05 and *p_crit_* = .01) and percentiles (97.5^th^ vs 99^th^) as threshold values; the t*_max_* correction using two thresholds (*p_crit_* = .05 and *p_crit_* = .01); and Bayesian t-tests with critical thresholds set at BF_10_ ≥ 3 and BF_10_ ≥ 10, plotted across a range of maximum Cohen’s *d* values from 0 to 2. (**B**) Percentage of electrodes surpassing the threshold across different statistical methods across Cohen’s d ranging from 0 to 2. **(C)** Percentage of datapoints and **(D)** electrodes that exceeded the thresholds within the range of Cohen’s *d* from 0 to 1 (highlighted by gray dashed squares in panels A and B).

#### Bayesian t-test

The Bayesian t-test started to detect timepoints and electrodes that exceeded the respective threshold of a BF_10_ ≥ 3 or BF_10_ ≥ 10 earlier than other methods, see Fig. 2. Notably, Bayesian t-tests identified condition differences below a small effect (i.e., below a Cohen’s d of *d* = .2). Specifically, from a maximum Cohen’s *d* of around *d* = .03, datapoints were flagged as exceeding the threshold (BF_10_ ≥ 3), suggesting that the test was sensitive to false positives, given that only small condition differences were simulated in the data. With a stricter threshold of BF_10_ ≥ 10, condition differences were detected from a Cohen’s *d* of *d* = .06, whereas other methods did not indicate any effects at this level. Relative to a threshold set at BF_10_ ≥ 3, a threshold of BF_10_ ≥ 10 detected fewer datapoints and indicated condition differences comparable to using a cluster-forming threshold of *p_crit_* = .01, see Fig. 2, and Fig. 3. The Bayesian approach was substantially more permissive than the *t*_max_ procedures at both thresholds of *p_crit_* = .05 and *p_crit_* = .01.

**Figure 3.**
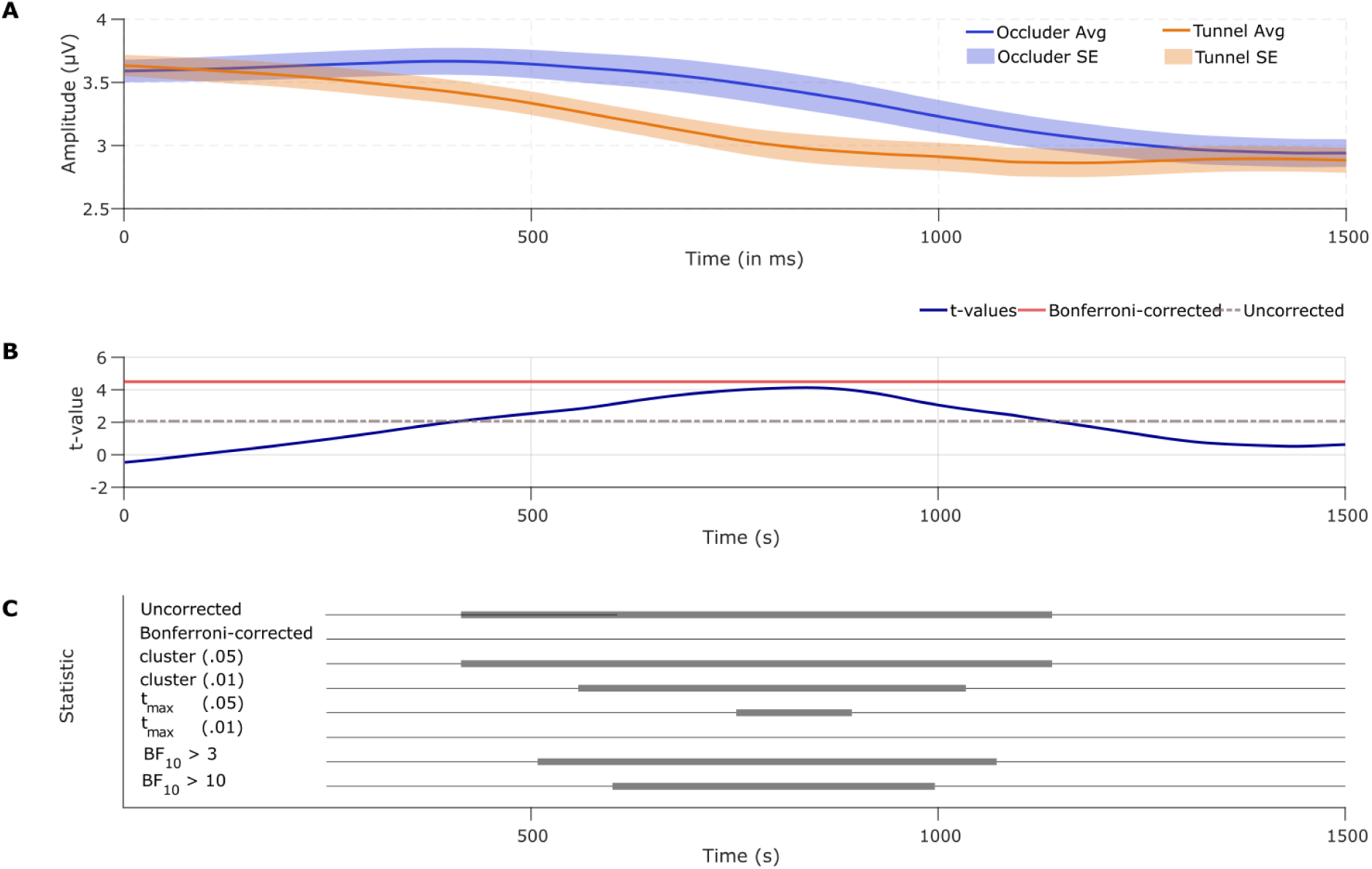
Statistical tests of steady-state evoked responses at a single electrode (Oz). (**A**) The evoked responses are shown, separately for the Occluder (blue) and the Tunnel condition (orange) for *N* = 40 participants. Shading indicates the standard error. (**B**) Time series of t-scores (blue) together with the Bonferroni-corrected (red) and uncorrected (grey, dashed) thresholds. (**C**) Significant timepoints for each of statistical procedures: uncorrected .05-level (two-sided), Bonferroni-corrected, the cluster-based permutation (forming-threshold at *p_crit_* = .05 and *p_crit_* = .01, no difference between the 97.5^th^ vs 99^th^ percentile), the t_max_ procedure (05-and .01-level), and a Bayes Factor threshold of BF_10_ ≥ 3 and BF_10_ ≥ 10. Note that the time series depicted for the cluster-based methods cannot be interpreted as point-wise significance, but only reflects cluster membership (see Sassenhagen & Draschkow, 2019).

### 3.2. Second Dataset: Event-Related Potentials

The simulation results for the dataset 2 are visualized in Fig. 3. Similar to dataset 1, increasing the statistical threshold, independent of technique applied, reduced the number of datapoints and electrodes considered statistically significant.

#### Cluster-based permutation

In dataset 2, the first cluster emerged at a small effect size of approximately *d* = .2. As the effect size increased, the number of electrodes contributing to the cluster gradually increased from below 5% to about half of all electrodes at *d* = 1.7 for a cluster-forming threshold at *p_crit_* = .05. The more conservative threshold (*p_crit_* = .01) resulted in fewer significant datapoints and electrodes included in clusters, showing comparable performance to a Bayes factor threshold of BF_10_ ≥ 10. Consistent with results from the first dataset, simulations for the ERP dataset demonstrated that the cluster-based permutation test identified the largest number of electrodes exceeding the significance threshold. This method showed the steepest increase in both significant timepoints and electrodes relative to the Bayesian t***-***test and the *t*_max_ procedure. Results observed when choosing the 97.5^th^ percentile in comparison to the 99^th^ percentile as cut-off were highly similar, also see Table 2.

#### T_max_ procedure

Corroborating simulations of the first dataset, the *t*_max_ permutation yielded the most conservative threshold compared to the other methods applied, also see Fig. 4 and Fig. 5. Only when a large effect size (*d* = .8) was simulated, the *t*_max_ procedure indicated small condition differences. Using a threshold of *p_crit_* = .05, the flagged datapoints using the *t*_max_ permutation at a large effect size corresponded to below 10 % of those detected with Bayesian t-tests (BF_10_ ≥ 3) or cluster-based permutation (*p_crit_* = .05). With a threshold of *p_crit_* = .01 for the *t*_max_ permutation, the detected datapoints decreased to around 5 % of those detected with Bayesian t-tests or the cluster-based permutation. With a Cohen’s *d* of 2, the *t*_max_ correction (*p_crit_* = .05) identified about 10% of all electrodes, corresponding to about 25% of the electrodes compared to those detected with a Bayesian t-test using a BF_10_ ≥ 3 and about 30% overlap with a Bayesian t-test threshold of BF_10_ ≥ 10.

**Figure 4.**
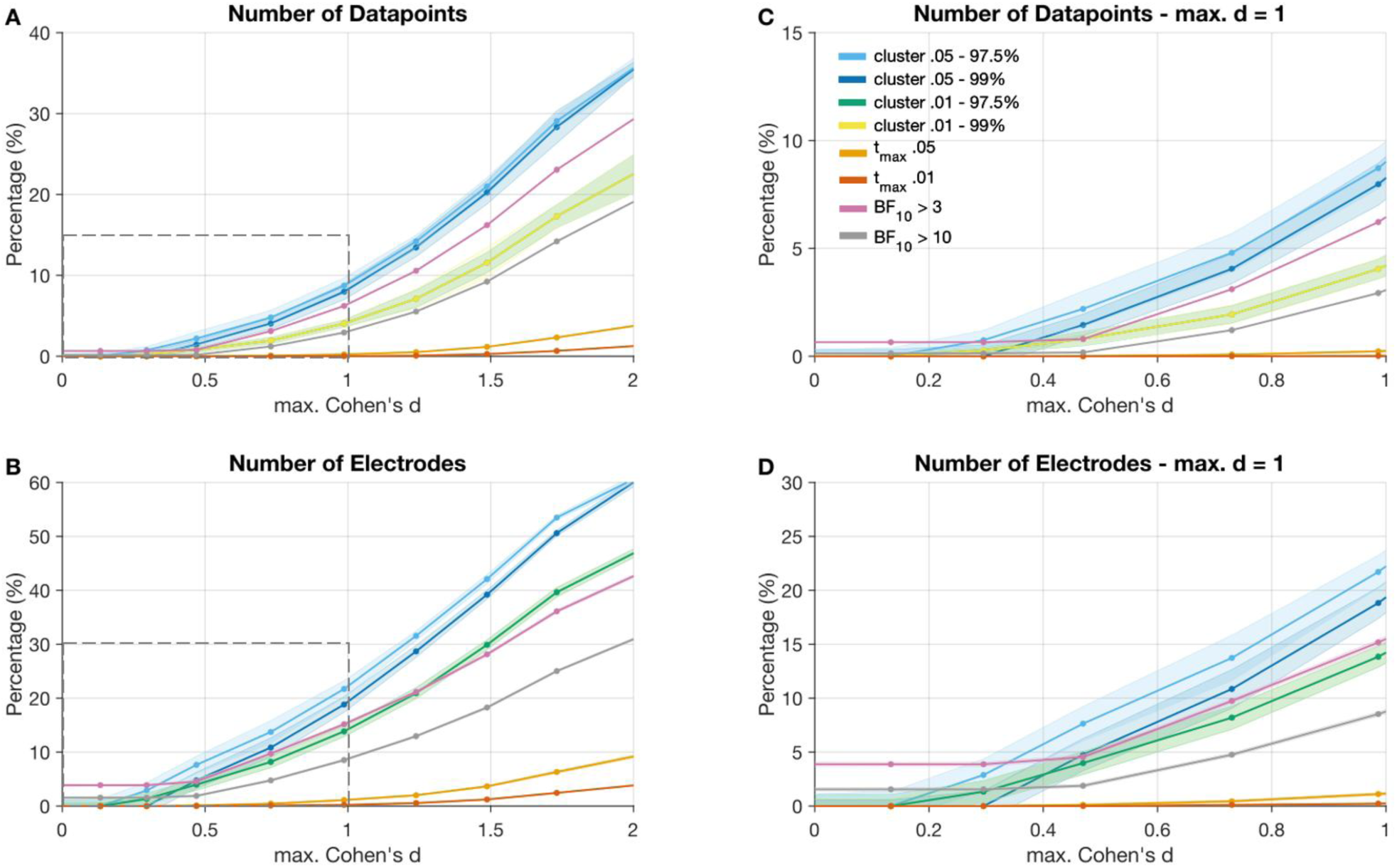
Results of the second dataset for simulated effect sizes for the ERP data after permutation. (**A**) Percentage of datapoints above the defined threshold for the respective mass univariate approach. (**B**) Percentage of electrodes exceeding the critical thresholds defined by the different statistical approaches: cluster-based permutation using different cluster-forming thresholds (*p_crit_* = .05 and *p_crit_* = .01) and percentiles (97.5^th^ vs 99^th^) as threshold values; the t*_max_* correction using different *t*-thresholds (*p_crit_* = .05 and *p_crit_* = .01); and Bayesian t-tests with thresholds set at BF_10_ ≥ 3 and BF_10_ ≥ 10, plotted across a range of maximum Cohen’s *d* values from 0 to 2. **(C)** Percentage of datapoints and **(D)** electrodes that exceeded the thresholds within the range of Cohen’s *d* from 0 to 1 (highlighted by gray dashed squares in panels A and B).

**Figure 5.**
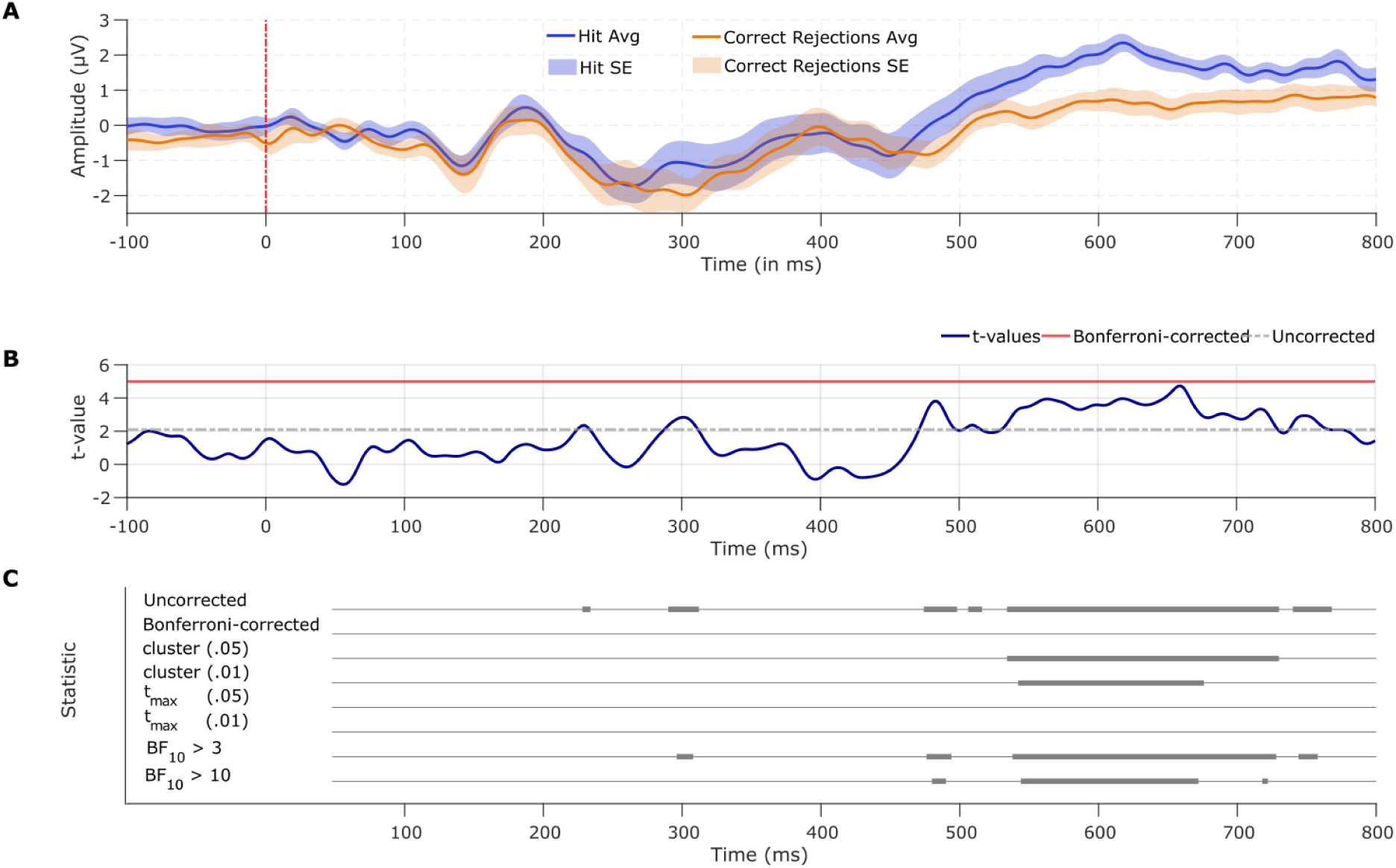
Statistical comparison of event-related potentials (old-new effect) at a single electrode (sensor 55). (**A**) Average evoked responses are shown, separately for hits (blue) and the correct rejections (orange) for *N* = 18 participants. Shading indicates the standard error. (**B**) Time series of *t-*scores (blue) and the Bonferroni-corrected (red) and uncorrected (grey, dashed) threshold. (**C**) Significant timepoints for each of statistical procedures: uncorrected .05-level (two-sided), Bonferroni-corrected, the cluster-based permutation threshold (forming-threshold at *p_crit_* = .05 and *p_crit_* = .01, no difference between the 97.5^th^ vs 99^th^ percentile), the t*_max_*procedure (05- and .01-level), and a Bayes Factor threshold of BF_10_ ≥ 3 and BF_10_ ≥10. Note that the time series depicted for the cluster-based methods cannot be interpreted as point-wise significance, but only reflects cluster membership (see Sassenhagen & Draschkow, 2019).

#### Bayesian t-test

The Bayesian approach detected condition differences at timepoints and electrodes at smaller effect sizes than other methods, i.e., before the cluster-based or *t_max_* methods indicated statistical differences. In line with the results of dataset 1, only Bayesian t-tests indicated condition differences at effect sizes below the conventional threshold for a small effect. Notably, datapoints greater than BF_10_ ≥ 3 were observed from a maximum Cohen’s *d* of approximately .03, despite only minimal condition differences. This pattern suggests that this approach likely yields false positives when a relatively small Bayes factor threshold is applied. Even the more conservative threshold of BF_10_ ≥ 10 flagged condition differences starting at around *d* = .03, whereas the other methods did not detect any effects at this level. Similar to the results of the first dataset, applying a threshold of BF_10_ ≥ 3 was highly similar to a cluster-forming threshold of *p_crit_* = .05, whereas applying a threshold of BF_10_ ≥ 10 revealed results comparable to a cluster-forming threshold of *p_crit_* = .01.

### 3.3. Third Dataset: Alpha Power

#### Cluster-based permutation

In dataset 3, the first cluster emerged at a small effect size below *d* = .2. As the effect size increased, the number of frequency bins contributing to the cluster gradually increased from around 10% of frequency bins when simulating a medium effect (*d* = .5). to around 30% of frequency bins at a large effect (Cohen’s *d* = .8) when choosing a cluster-based permutation with a forming threshold of *p_crit_* = .05. Result patterns were highly similar for the 97.5^th^ percentile cutoff and the 99^th^ percentile cutoff, also see Table 3. As expected, when a stricter cluster forming thresholds was applied (.01), the cluster-based permutation was a bit more conservative.

#### T_max_ procedure

In line with simulation results of dataset 1 and 2, the *t*_max_ permutation yielded the most conservative threshold compared to the other methods applied. With a *p_crit_* = .05, the *t*_max_ procedure flagged condition differences for small effects (*d* = .2). Using a critical threshold at *p_crit_* = .01, only effects with a medium effect size were detected. With a large effect (Cohen’s *d* = .8), the *t*_max_ permutation indicated condition differences in approximately half of the datapoints and around half of the frequency bins compared to the cluster-based permutation (both *p_crit_* = .05). Approximately 50% of the datapoints and 25% of the frequency bins were considered significant under the *t*_max_ compared to the cluster-based permutation with Cohen’s *d* reaching 1.5, see Fig. 6.

**Figure 6.**
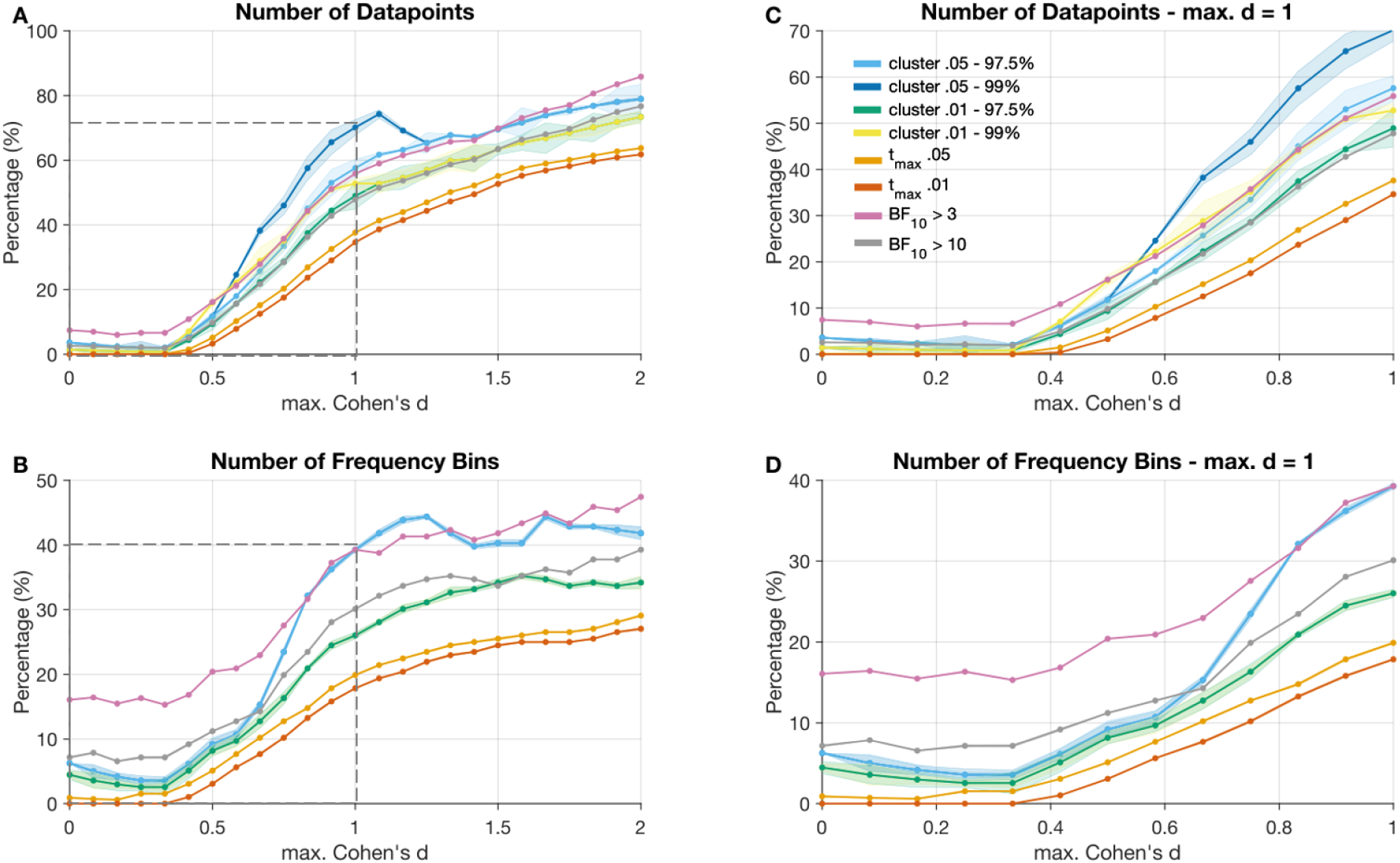
Results of the third dataset for simulated effect sizes for the alpha power reduction. (**A**) Percentage of datapoints above the defined threshold for the respective mass univariate approach. (**B**) Percentage of frequency bins exceeding the critical thresholds defined by the different statistical approaches: cluster-based permutation using different cluster-forming thresholds (*p_crit_* = .05 and *p_crit_* = .01) and percentiles (97.5^th^ vs 99^th^) as threshold values; the t*_max_* correction using different *t*-thresholds (*p_crit_* = .05 and *p_crit_* = .01); and Bayesian t-tests with thresholds set at BF_10_ ≥ 3 and BF_10_ ≥ 10, plotted across a range of maximum Cohen’s *d* values from 0 to 2. **(C)** Percentage of datapoints and **(D)** frequency bins that exceeded the thresholds within the range of Cohen’s *d* from 0 to 1 (highlighted by gray dashed squares in panels A and B).

#### Bayesian t-test

The Bayesian t-test with a threshold of BF_10_ ≥ 3 and BF_10_ ≥10 performed comparable to the cluster-based permutation approach with a cluster forming threshold of *p_crit_* = .05 and *p_crit_* = .01, respectively. Critically, the Bayes Factor indicated condition differences in timepoints and electrodes earlier than the cluster-based or *t_max_* method. As described for dataset 1 and 2, the Bayes Factor identified condition differences when small effects were simulated (Cohen’s *d* < .2, see Fig. 6). Further, the Bayesian t-test detected condition differences from the beginning, where no condition differences were simulated (false positives).

## 4. Discussion

The present study compared different mass univariate statistical approaches that aim to control for the multiple comparison problem, using real and simulated EEG data. Mass univariate analyses involve a large number of comparisons with corrections for the increased probability of false positive findings. The outcomes and results greatly depend on the procedure used to control for multiple comparisons. Here, we contrast cluster-based permutation, the t*_max_* permutation, and Bayesian t-tests across a range of simulated effects size in the time- and frequency domain, in three EEG datasets.

To explore the implications of different methods, we compared the amount of datapoints and electrodes or frequency bins shown as statistically meaningful when simulating condition differences of increasing effect size. Overall, the choice of the mass univariate approach, i.e., whether t_max_, cluster-based permutation, or Bayes Factor thresholds were used, had a substantial impact on the number of datapoints (timepoints by electrodes or timepoints by frequency bins) identified as crossing a given threshold. Our findings suggest that the cluster-based permutation approaches and Bayesian t-tests provided a more liberal approach and yielded more broadly distributed clusters, whereas the *t*_max_ procedure was most conservative and indicated more spatially restricted and focal clusters.

### 4.1. Cluster-based permutation

As demonstrated in the simulations, the sensitivity of the cluster-based permutation approach varied as a function of spatial sensor layout. As detailed in prior work, the respective cluster-mass statistic depends on the number of electrodes included in the layout as well as the neighborhood definition and if a minimum number of cluster members is defined a priori (also see (Groppe et al., 2011a). In addition, the results vary depending on the cluster-forming threshold used for cluster inclusion. Using more extreme thresholds has been shown to result in more distinct, focal clusters, making the procedure particularly effective for detecting broad effects (Groppe et al., 2011b). Cluster-based permutation highlights ERP effects or effects with temporal and spatial continuity because the method captures the coherent patterns typical of ERPs, which are less likely to arise from random noise. However, the same principle makes the method less sensitive to highly localized or focal effects that consist of only a few time points or electrodes. In line with this, our simulations suggested that the procedure might overlook effects that are narrowly distributed or very short, as illustrated by the fact that the approach first started to indicate condition differences when effect sizes were large. In addition, it demonstrates that the test might miss significant effects when those are part of a bigger cluster with very high *t*-scores nearby. Under certain conditions, the method might also merge distinct ERP effects in the same cluster as broadly distributed effects, spanning many time points and electrodes (see Groppe et al., 2011 for simulations of the P2 and P3 component). In contrast, other studies have shown that when statistical power is low, cluster-based methods may include a substantial number of false positives, potentially leading to spurious interpretations of the data (Fields & Kuperberg, 2020). Finally, it is important to note that different parameters will considerably affect the test’s outcome (e.g., the cluster-inclusion to form a cluster or the percentile used to define the significance threshold of the null distribution; Groppe et al., 2011; Maris & Oostenveld, 2007; Hemmelmann et al, 2004). However, the current simulations suggest that choosing a stricter percentile to determine a cluster’s statistical significance (e.g., 97.5^th^ vs. 99^th^) has little impact on the resulting cluster extent. Future research may wish to examine the impact of the choice of the specific test statistic (e.g., *t*_min_, maximum size of a cluster, maximum *t*-score) on the results.

### 4.2. t_max_

In contrast to cluster-based methods, our simulations suggest that the *t*_max_ permutation is particularly sensitive to effects that are more narrowly distributed. Simulations showed that the resulting time point by electrode matrices exceeding the threshold defined via t*_max_* permutation were the most conservative among the methods compared in the present study. While the technique provides strong control of the FWER (Groppe et al., 2011b), it leads to fewer datapoints that are considered statistically significant, rendering the *t*_max_ procedure the most conservative test among the contrasted techniques.

### 4.3. Bayesian methods

Across a range of simulations assessing the methods’ permissiveness, Bayesian t-tests were more liberal than the *t*_max_ statistic but more conservative than cluster-based permutation methods, even those with more extreme thresholds (such as a *p-*value of *p_crit_* = .01). In scenarios where strong a-priori time windows and spatial regions are utilized as priors, the Bayesian t-test may prove especially advantageous. By incorporating prior knowledge into the model, this method can enhance the precision of statistical inference and allow for a more targeted, hypothesis-driven, analysis compared to traditional methods like time window averaging. In addition, the Bayesian approach offers a continuous measure of evidence for or against an effect. Specifically, it quantifies how likely it is that there is an effect over the likelihood of no effect, thus facilitating model comparisons and quantifying the uncertainty given the data. However, these tests might be overly sensitive to deviations in variance between populations or differences in sample size, which may be misattributed as condition differences. Particularly, Bayesian t-tests were consistently indicating meaningful condition differences when the maximum effect size was reported to be lower than *d* = .3, independent of the dataset. That is, Bayesian t-tests indicated condition differences in electrodes or frequency bins exceeding a given threshold when other methods did not support such conclusions at effect sizes that were neglectable. Both the *t*_max_ permutation and the Bayesian t-test do not require clusters of connected points across space, time, or frequency to exceed a predefined threshold. In contrast, each point is treated independently (although EEG data is highly correlated). As a result, in the present simulation, the Bayesian t-test was sensitive to differences that might have resulted from noisy or very short-lived condition differences, which may represent false positives rather than meaningful effects. For these cases, a stricter threshold or bootstrapping approaches in addition to careful prior modeling might be more appropriate (Ahumada et al., 2025; Rousselet, 2025; Schwarzkopf, 2015).

### 4.4. Conclusion

Mass-univariate analyses provide a flexible method for examining the extent of temporal and spatial characteristics of condition differences in studies using human EEG recordings. Bayesian statistics offer a highly valuable and easy-to-implement alternative to the more traditional frequentist (NHT) approaches used to determine the existence of an effect, and its temporal and spatial extent. Mass-univariate analyses often yield greater statistical power compared to traditional time window averaging methods, which may dilute effects by including irrelevant time points or spatial locations. By incorporating prior knowledge about the expected timing and localization of effects into mass-univariate approaches, researchers can reduce noise and improve the precision of their inferences.

## Data availability

All data is available on OSF (https://osf.io/8vs42/?view_only=ea96f065cd774e0992ade6f3a58e4d53).

## Acknowledgments

This work has been supported by: Grant R01MH125615 from the National Institutes of Health; National Science Foundation grant number 2318984.

## Author contributions: CRediT

- Conceptualization: AK
- Data curation: AT, AK, CP
- Formal analysis: AT, AK, CP
- Funding acquisition: AK
- Investigation: AT, AK, CP
- Methodology: AK, AT, CP
- Project administration: AK, AT
- Resources: AK, AT
- Supervision: AK, AT
- Visualization: AT
- Writing – original draft: AT
- Writing – review and editing: AK, AT, CP

