## Supplemental Tables for "Comparing Methods for Mass Univariate Analyses of Human EEG: Empirical Data and Simulations"

**Dataset 1: SSVEP (*N*=40)**

| | Threshold (percentile) | Datapoints (%) | | | | Electrodes (%) | | | |
|---|---|---|---|---|---|---|---|---|---|
| | | *d* = 0 | *d* = .2 | *d* = .5 | *d* = .8 | *d*=0 | *d*=.2 | *d*=.5 | *d*=.8 |
| cluster-based | .05 (97.5th) | 0 | 0 | 13.2 | 43.4 | 0 | 0 | 59.3 | 89.0 |
| | .05 (99th) | 0 | 0 | 12.2 | 43.4 | 0 | 0 | 54.4 | 89.0 |
| | .01 (97.5th) | 0 | 0.3 | 5.0 | 25.4 | 0 | 1.6 | 26.9 | 78.6 |
| | .01 (99th) | 0 | 0 | 4.6 | 24.6 | 0 | 0 | 25.3 | 78.6 |
| $t_{max}$ | .05 | 0 | 0 | 1.0 | 6.9 | 0 | 0 | 6.6 | 31.3 |
| | .01 | 0 | 0 | 0.3 | 2.3 | 0 | 0 | 2.7 | 12.1 |
| $BF_{10}$ | 3 | 0.5 | 1.8 | 7.6 | 33.6 | 5.8 | 9.9 | 37.4 | 79.1 |
| | 10 | 0 | 0.6 | 3.5 | 20.6 | 0 | 4.9 | 18.1 | 68.1 |

*Table 1.* Percentage of datapoints and electrodes detected under various statistical methods (cluster-based permutation, $t_{max}$ permutation, Bayesian t-test) using different thresholds across different effect sizes (Cohen's *d* = 0, .2, .5, .8, corresponding to no, small, medium, and large effects) in the first dataset (SSVEP responses).

**Dataset 2: Event-Related Potential (old-new effect, *N*=18)**

| | | Datapoints (%) | | | | Electrodes (%) | | | |
|---|---|---|---|---|---|---|---|---|---|
| | Threshold (percentile) | *d* = 0 | *d* = .2 | *d* = .5 | *d* = .8 | *d*=0 | *d*=.2 | *d*=.5 | *d*=.8 |
| cluster-based | .05 (97.5th) | 0 | 0 | 2.2 | 4.8 | 0 | 0 | 7.6 | 13.7 |
| | .05 (99th) | 0 | 0 | 1.5 | 4.1 | 0 | 0 | 4.8 | 10.9 |
| | .01 (97.5th) | 0 | 0 | 0.8 | 1.9 | 0 | 0 | 4.0 | 8.2 |
| | .01 (99th) | 0 | 0 | 0.8 | 1.9 | 0 | 0 | 4.0 | 8.2 |
| $t_{max}$ | .05 | 0 | 0 | 0.01 | 0.1 | 0 | 0 | 0.1 | .4 |
| | .01 | 0 | 0 | 0 | 0 | 0 | 0 | 0 | .1 |
| $BF_{10}$ | 3 | 0.7 | 0.7 | 0.8 | 3 | 3.9 | 3.9 | 4.5 | 9.7 |
| | 10 | 0.1 | 0.1 | 0.2 | 1.2 | 1.6 | 1.6 | 1.9 | 4.8 |

*Table 2.* Percentage of datapoints and electrodes detected under various statistical methods (cluster-based permutation, $t_{max}$ permutation, Bayesian t-test) using different thresholds across different effect sizes (Cohen's *d* = 0, .2, .5, .8, corresponding to no, small, medium, and large effects) in the second dataset (ERP responses to old-new effect).

**Dataset 3: Alpha Power (*N*=50)**

| | | Datapoints | | | | Electrodes (max. 129) | | | |
|---|---|---|---|---|---|---|---|---|---|
| | Threshold (percentile) | *d = 0* | *d = .2* | *d = .5* | *d = .8* | *d=0* | *d=.2* | *d=.5* | *d=.8* |
| cluster-based | .05 (97.5th) | 3.6 | 2.1 | 18 | 45 | 6.3 | 3.6 | 10.7 | 32.1 |
| | .05 (99th) | 3.6 | 2.1 | 24.6 | 57.6 | 6.3 | 3.6 | 10.7 | 32.1 |
| | .01 (97.5th) | 1.5 | 0.83 | 15.6 | 37.5 | 4.5 | 2.6 | 9.7 | 20.9 |
| | .01 (99th) | 1.5 | 0.83 | 22.2 | 44 | 4.5 | 2.6 | 9.7 | 20.9 |
| $t_{max}$ | .05 | 0.1 | 0.12 | 10.3 | 26.9 | 0.9 | 1.5 | 7.7 | 14.8 |
| | .01 | 0 | 0 | 7.9 | 23.7 | 0 | 0 | 5.6 | 13.3 |
| $BF_{10}$ | 3 | 7.5 | 6.6 | 21.2 | 44.3 | 16.1 | 16.3 | 20.9 | 31.6 |
| | 10 | 2.6 | 2.2 | 15.6 | 36.3 | 7.1 | 7.1 | 12.8 | 23.5 |

*Table 3.* Percentage of datapoints and electrodes detected under various statistical methods (cluster-based permutation, $t_{max}$ permutation, Bayesian t-test) using different thresholds across different effect sizes (Cohen's *d* = 0, .2, .5, .8, corresponding to no, small, medium, and large effects) in the third dataset (alpha power differences).
